# Genetic Fingerprinting of Herbarium Hop Specimens Reveals Genetic Diversity and Relationships with Modern Finnish Hops

**DOI:** 10.1101/2025.11.12.687964

**Authors:** Thien-Tam Luong, Teija Tenhola-Roininen, Merja Hartikainen, Mia Lempiäinen-Avci, Sanna Huttunen, Lidija Bitz

## Abstract

We sampled 351 hop herbarium specimens for genetic analyses of their identities through definition of unique set of samples by extraction of duplicates and comparative analyses with modern hop samples from Finland and worldwide. We also analyzed genetic diversity and assessed genetic clustering and structure within and between herbarium hops specimens and modern hop germplasm. A set of 26 previously published hop microsatellites (SSRs) proven in diversity analyses have been used for genetic evaluations of herbarium hop DNA extracted from pressed and dried specimens of which the oldest was dated back to the year of 1841. A genetic fingerprint was generated for all samples and unique ones has been generated in pairwise comparisons of pairs of samples and eliminations of duplicates. For unique set of samples, a genetic distance matrix was created for clustering analyses as well allele frequencies matrix for model-based clustering analyses. Herbarium hop samples representing historical DNA showed no clear differentiation (clustering or sub-clustering) when analyzed with presently grown modern hops. Weak tendency of hops to cluster according to the geographical origin has been noticed. We detected 39 groups of hop herbarium duplicates representing 94 specimens, where redundancy showed to be low when compared to modern Finnish hops (redundancy >70%). Analyzing herbarium hops through DNA extraction will add another link in investigating the origin and spread of Finnish hops from ancient times. Finnish hops has been described as a millennia old and native to Finland.

## Introduction

Native hop (*Humulus lupulus*) reaches its northernmost range in Finland (66°19′N), just 26 km south of the Arctic Circle (Suominen, 1994), and has recently been recorded even farther north near Inari (68°54′; ETRS89) (Lampinen & Lahti, 2018). While mainly found in southern Finland, occasional northern populations grow in nutrient-rich riverine woods. Hop is dioecious, often forming single-sex stands along stream rapids, and its Finnish name (*humala*) appears in many local waterway names.

Pollen evidence from Lake Huhdasjärvi shows hop presence from 8500 BC to 1195 AD—millennia before cultivation began (Alenius et al., 2013). Seeds from 11th–13th century settlements confirm its use before Finland, under Swedish rule (1200–1808), began taxing hops. By the 15th–16th centuries, hops were widely cultivated; King Kristoffer’s law (1442) required 40 hop poles per farm, and Turku paid tithes in hops while also importing cones from Gdansk (Leskelä, 2010). Brewing records from the 1540s mention beers with varying hop content, and by the 17th century hop yards were common—Laukko Manor reported 280 kg in 1699.

Beyond brewing, hops served as medicine, flavoring for *sima*, vegetables, hair dye, mattress stuffing, and fibers for ropes and fabrics (Ruoff, 2001). Cultivation declined in the 19th century as imported hops replaced local varieties. Interest revived about a decade ago with the rise of microbreweries (now one per ∼55,000 inhabitants). Today, hops are valued not only for brewing but also for their phytoestrogens, flavonoids, and fibers, driving efforts to select and cultivate the best local clones for future production.

## Materials and methods

### Plant materials

#### Herbarium collection survey and data

A set of 351 hop herbarium samples were sampled from herbarium department at University of Turku (Finland).

#### Modern hop samples

A set of containing >1000 hops contemporary growing in Finland has been previously genotyped and added to the analyses of this study.

### DNA extractions, microsatellite (SSR) selection and genotyping

A total DNA was extracted from herbarized specimens by EZNA/Qiagen Plant kit. An amount of 10 ng of DNA was used for multiplexing PCR reaction in which 3 or 4 differently dyed microsatellites have been combined (Supplemental table I)

A set of 26 microsatellite markers was selected from previous studies (Jakše et al., 2008; Jakše et al., 2011; Brady et al., 1996; Stajner et al., 2005; Hadonou et al., 2004), considering the species from which they were developed, their polymorphism information content (PIC), functionality, degree of polymorphism, and ease of interpretation (Supplementary Table I). For fragment analysis, one primer of each pair was labeled with a fluorescent dye—FAM™ (5-carboxyfluorescein), NED™, VIC®, or PET®—to enable separation and visualization of amplification products using an ABI PRISM® 310 Genetic Analyzer (Thermo Fisher Scientific Inc., Vantaa, Finland). PCR amplification was performed in 10 µl reactions containing 5 µl Thermo Scientific Phusion U Multiplex PCR Master Mix (Thermo Fisher Scientific Inc.), 20 ng of extracted hazelnut DNA, and 200–400 nM of each primer. The 26 SSR markers were amplified in two multiplex PCR reactions. Multiplex design was optimized using Multiplex Manager v1.2 (http://multiplexmanager.com) (Table 2), with the first five markers amplified in one reaction at an annealing temperature of 58 °C and the remaining four in a second reaction at 54 °C. PCR cycling was carried out on a BioRad C1000 Thermal Cycler (Bio-Rad, Hercules, California, USA) under the following conditions: 40 cycles of 10 s at 98 °C, 30 s at the respective annealing temperature, and 30 s at 72 °C, preceded by an initial denaturation at 98 °C for 30 s and followed by a final extension at 72 °C for 5 min. PCR products were diluted 1:50 prior to capillary electrophoresis on the ABI platform, and allele sizes were estimated using GeneMapper Software v5.

#### Detection of Duplicates

Each pair of genotypes, herbarium specimens, or samples was compared against all others, and those exhibiting identical SSR profiles across all analyzed loci (allowing for up to two mismatches) were grouped as duplicates (Supplemental Data II). These duplicate entries were removed from subsequent analyses, and one representative from each duplicate group was retained and incorporated into the unique sample set for further genetic clustering and diversity assessments.

#### Genetic clustering analyses

Genetic diversity parameters—including the number of alleles, observed heterozygosity (Ho), expected heterozygosity (He), and polymorphic information content (PIC)—were calculated for each marker using CERVUS version 3.0.7 (Kalinowski et al., 2007).

Distance-based clustering was performed in DARwin version 6.0.015 (Perrier et al., 2003) by computing the simple matching dissimilarity coefficient and constructing a dendrogram using the unweighted neighbor-joining method based on these distance measures.

Population structure was investigated using STRUCTURE version 2.3.4 with admixture model and correlated allele frequencies (Pritchard et al. 2000; Falush et al. 2003; Hubisz et al. 2009). A burn-in period of 200,000 interactions and 1,000,000 Markov Chain Monte Carlo (MCMC) iterations were used. The most probable number of genetic groups (K) was evaluated according to Evanno et al. (2005) in STRUCTURE HARVESTER (Earl and von Holdt 2012) by running 10 times for each K value from 1 to 23 (Evano et al. 2005). In order to maximize similarity on multiple STRUCTURE runs (10) for each single K value, membership coefficient matrices (Q-matrices) were permuted to have the closest match by applying Large K Greedy algorithm from the program CLUMPP version 1.1.2 (Jakobsson and Rosenberg 2007). The mean of the permuted Q-matrices was then used in the program DISTRUCT version 1.1 (Rosenberg 2004) to generate an output for visual representation of the aligned cluster assignments.

The program GenAlEx 6.5 (Peakall and Smouse 2006, 2012) was used for counting genetic diversity indices for clusters: He, Ho, Shannon’s diversity index I (Shannon 1948), and the percentage of polymorphic loci. Significance of the results was tested by permuting the SSR marker data 999 times. Genetic distances between individuals were estimated and an unweighted neighbor-joining (NJ, Saitou and Nei 1987) radial tree based on simple matching dissimilarity matrix was built with Darwin 6.0.014 (Perrier et al. 2003) with a bootstrap value of 1,000.

To visualize dissimilarities between clusters, principal coordinates analysis (PCoA) based on Nei’s genetic distances (Nei 1972) was performed with GenAlEx 6.5. Correlation between genetic distances (Nei’s distance) with geographic distances (km) of populations was tested with a Mantel test (Mantel 1967) in the software GenAlEx. Significance was assessed by conducting 999 permutations.

## Results

### Groups of duplicates

Duplicate analyses revealed 39 groups of duplicates containing two till eight specimens (Supplemental data II). Have they been accidently sampled from very same plant individuals or is there a special reason why particularly those specimens were clonally propagated and dispersed around.

<u>Genetic clustering</u> representing herbarium hop specimens only (Figure 1 and Supplemental data III)

**Figure 1.**
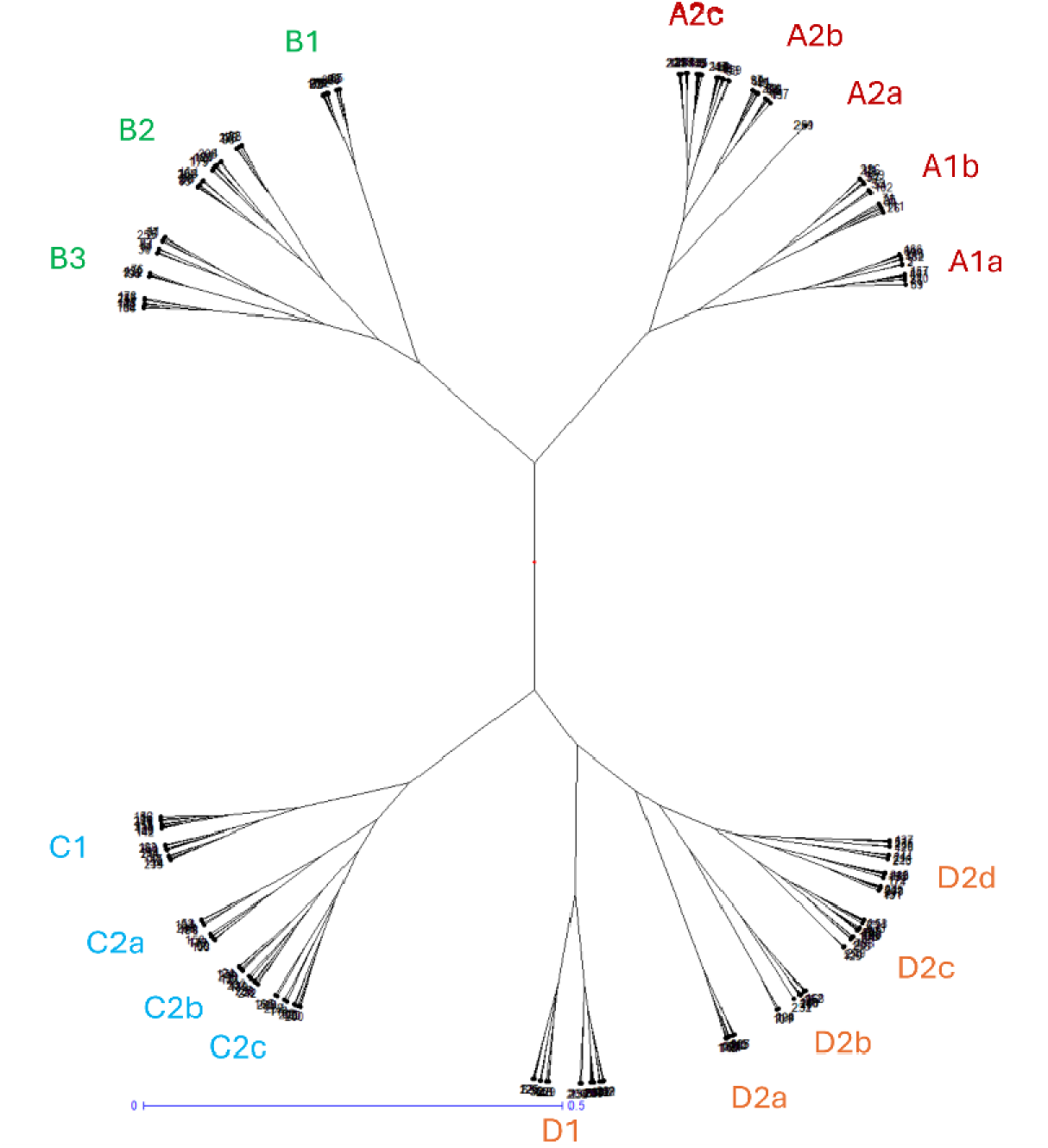
Genetic dendrogram representing unique herbarium hop specimens (257) samples.

**Figure 2.**
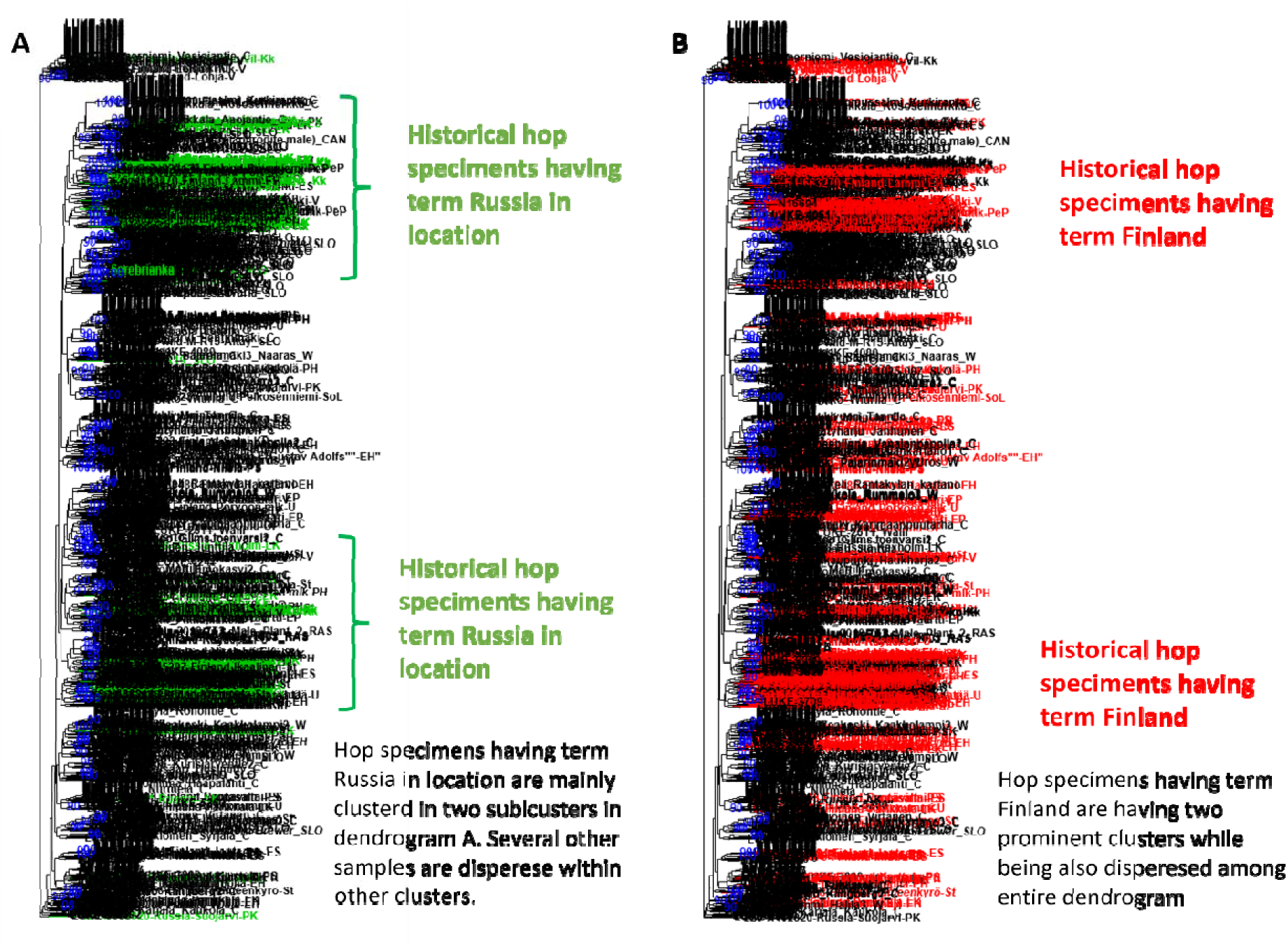
Genetic clustering analyses of >1500 hop genotypes including >300 herbarium hop specimens using 10 microsatellites (SSRs) A Dendrogram where only samples originating from Russia are labeled. B Dendrogram with labeled samples from Finland. Samples from Russia seems to tend to group into two clusters on the dendrogram as shown on A. while Finland samples are dispersed across the dendrogram. It must be mentioned that there is more Finland samples what might bias clustering.

---

**Supplemental table I.**
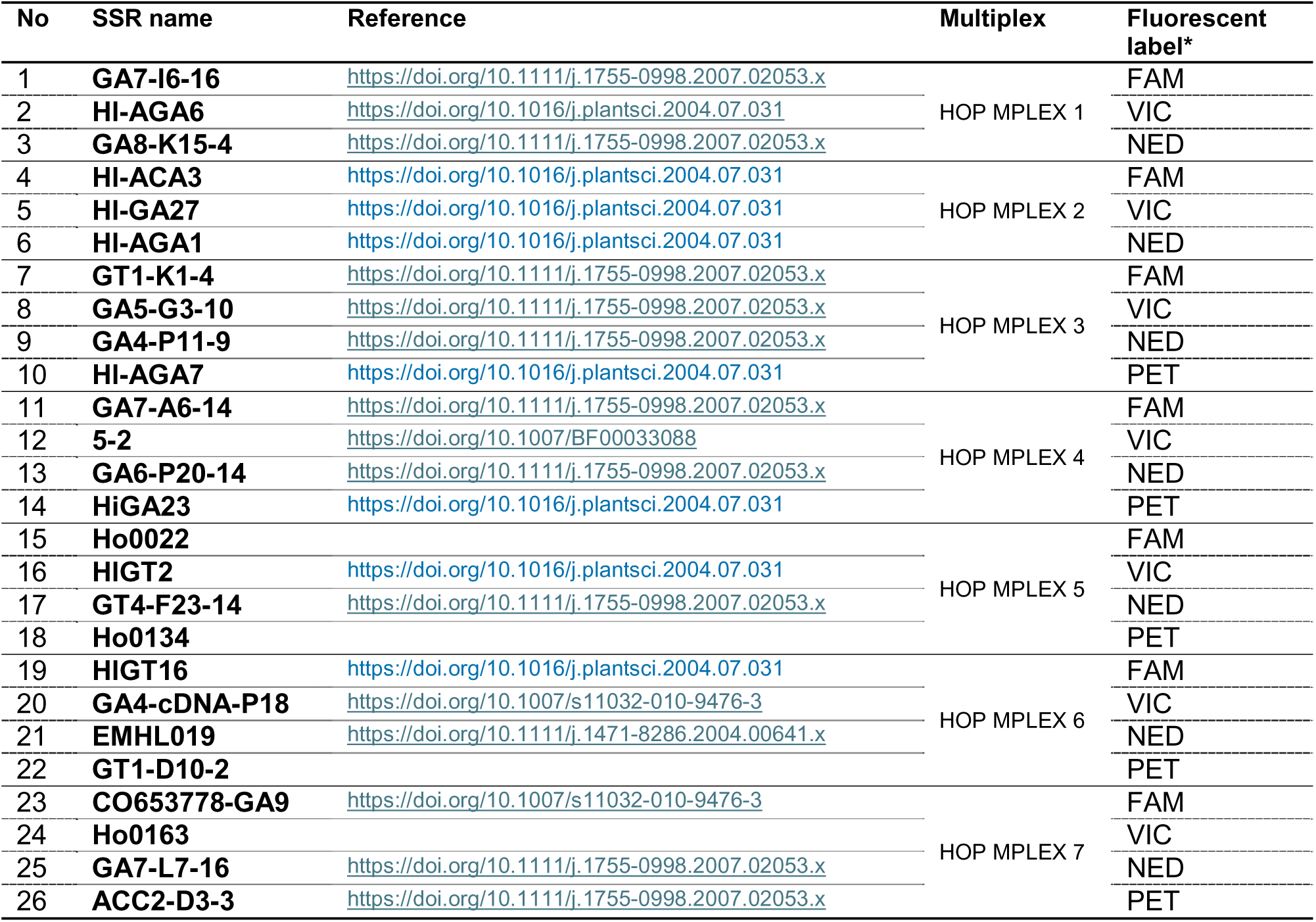

## Supplemental data II

Groups of hop duplicates and list of samples of each duplicate group with extended information on country of origin, locality name, biogeographical region and habitat description

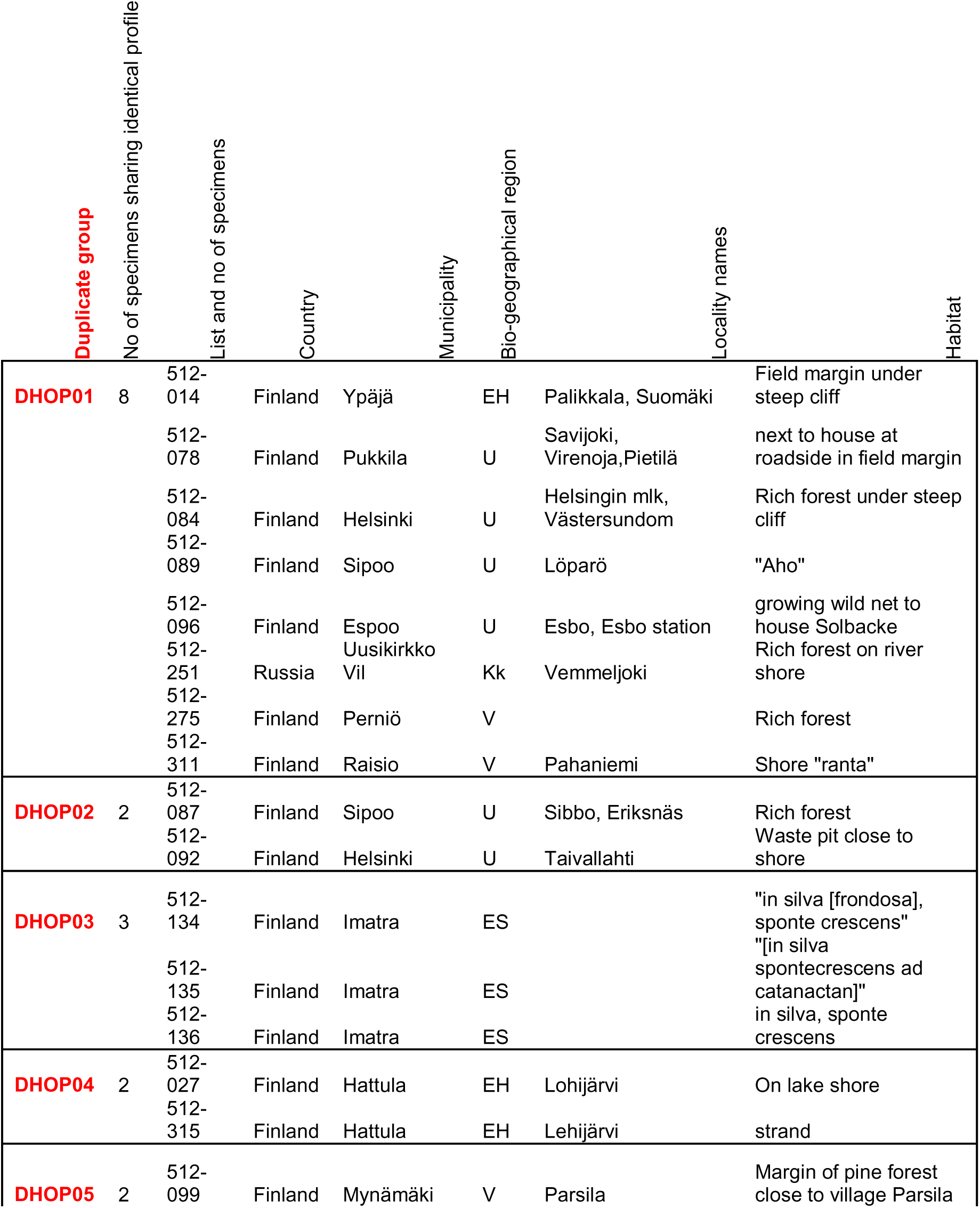
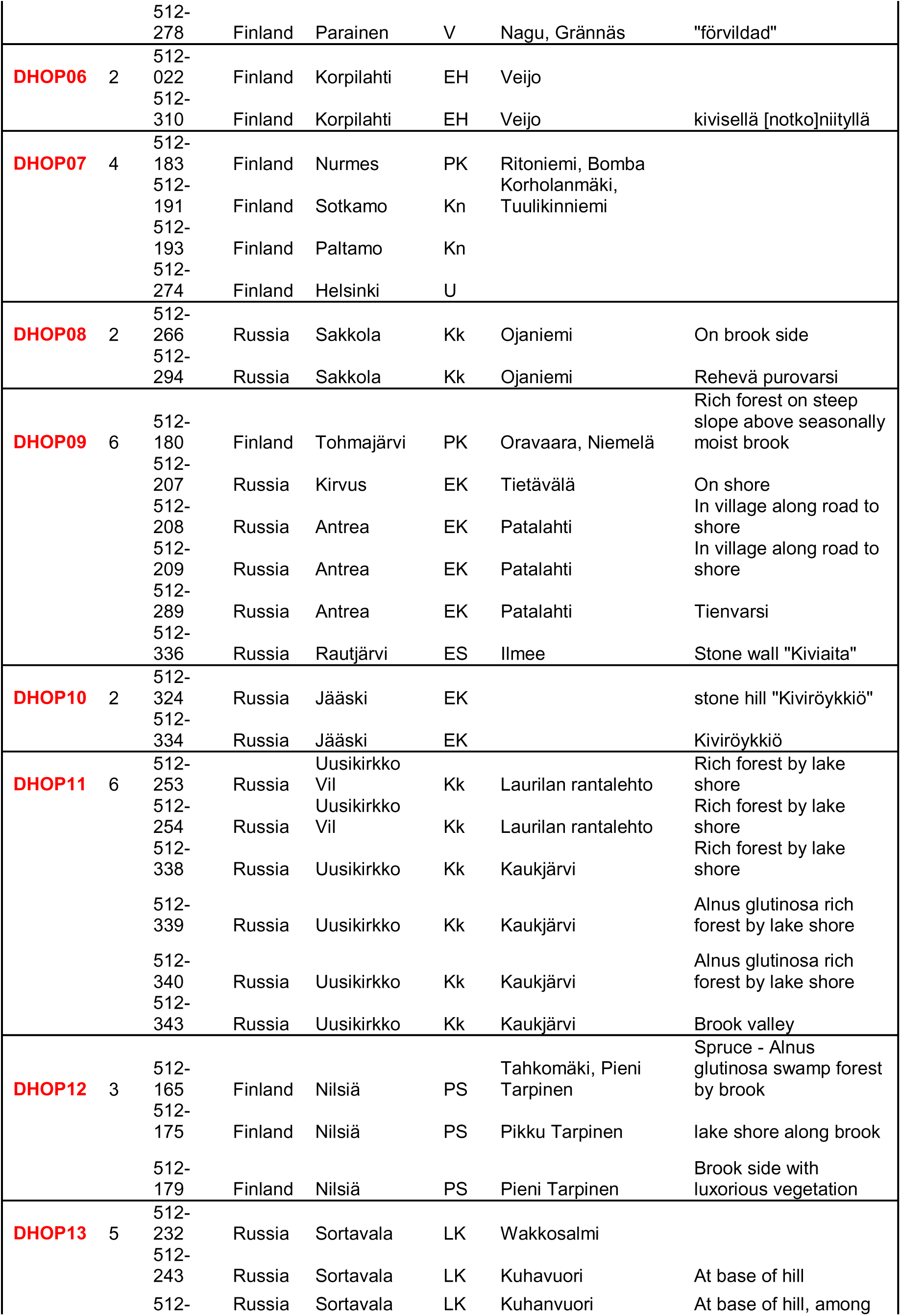
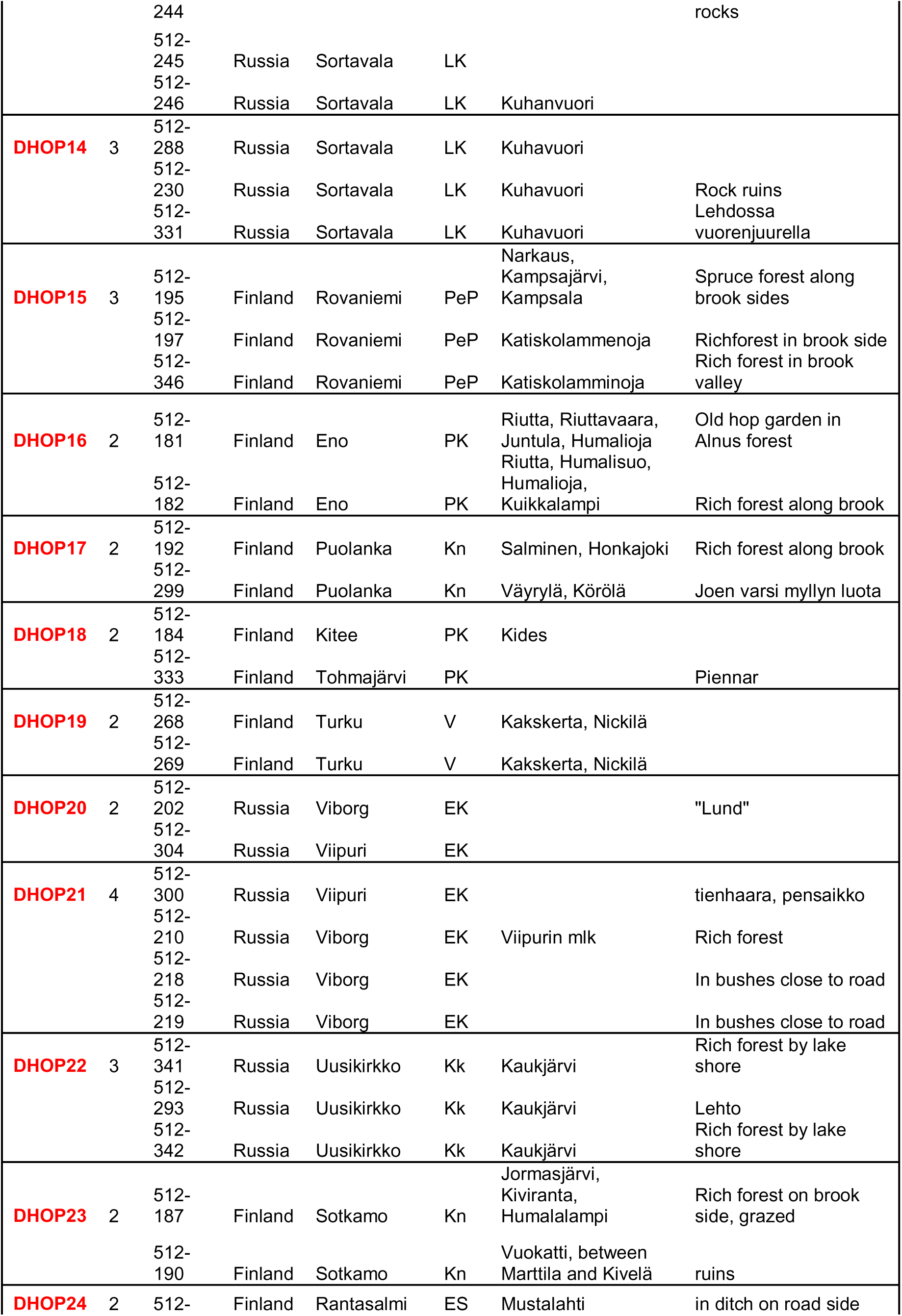
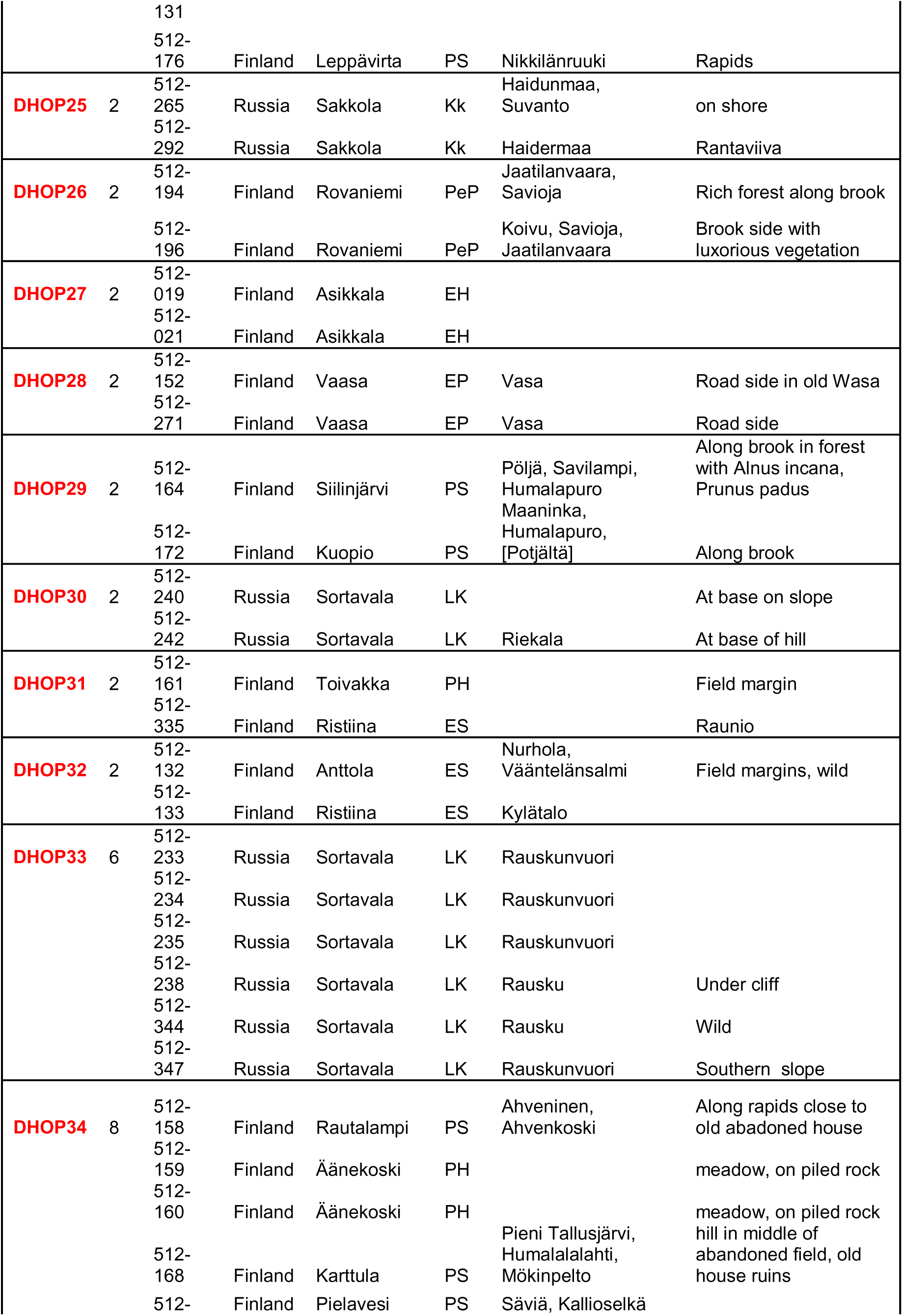
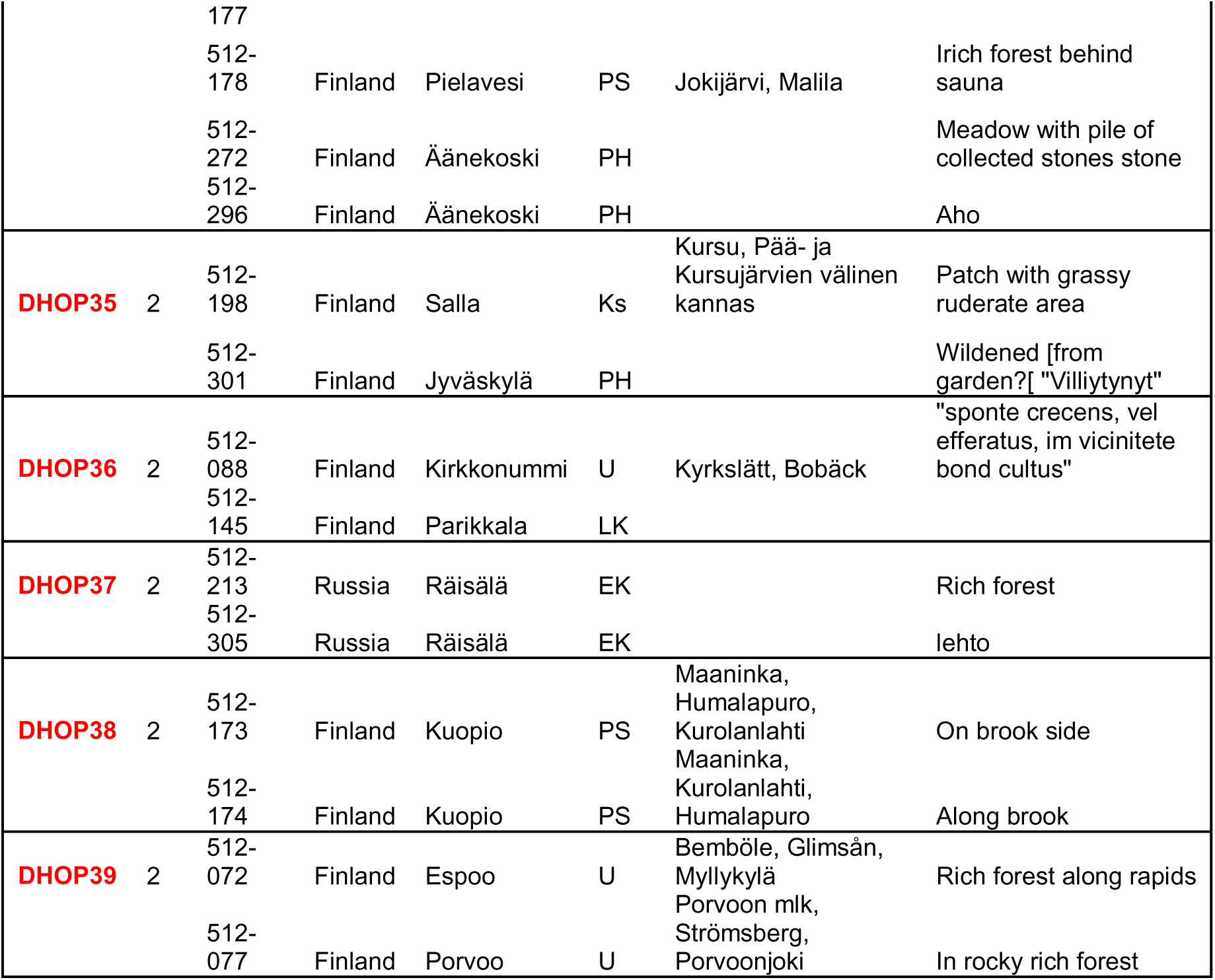

## Supplemental data III

Genetic clusters of herbarium hops based on hierarchical clustering (Ward coeff) and neighbor joining (simple matching coeff)

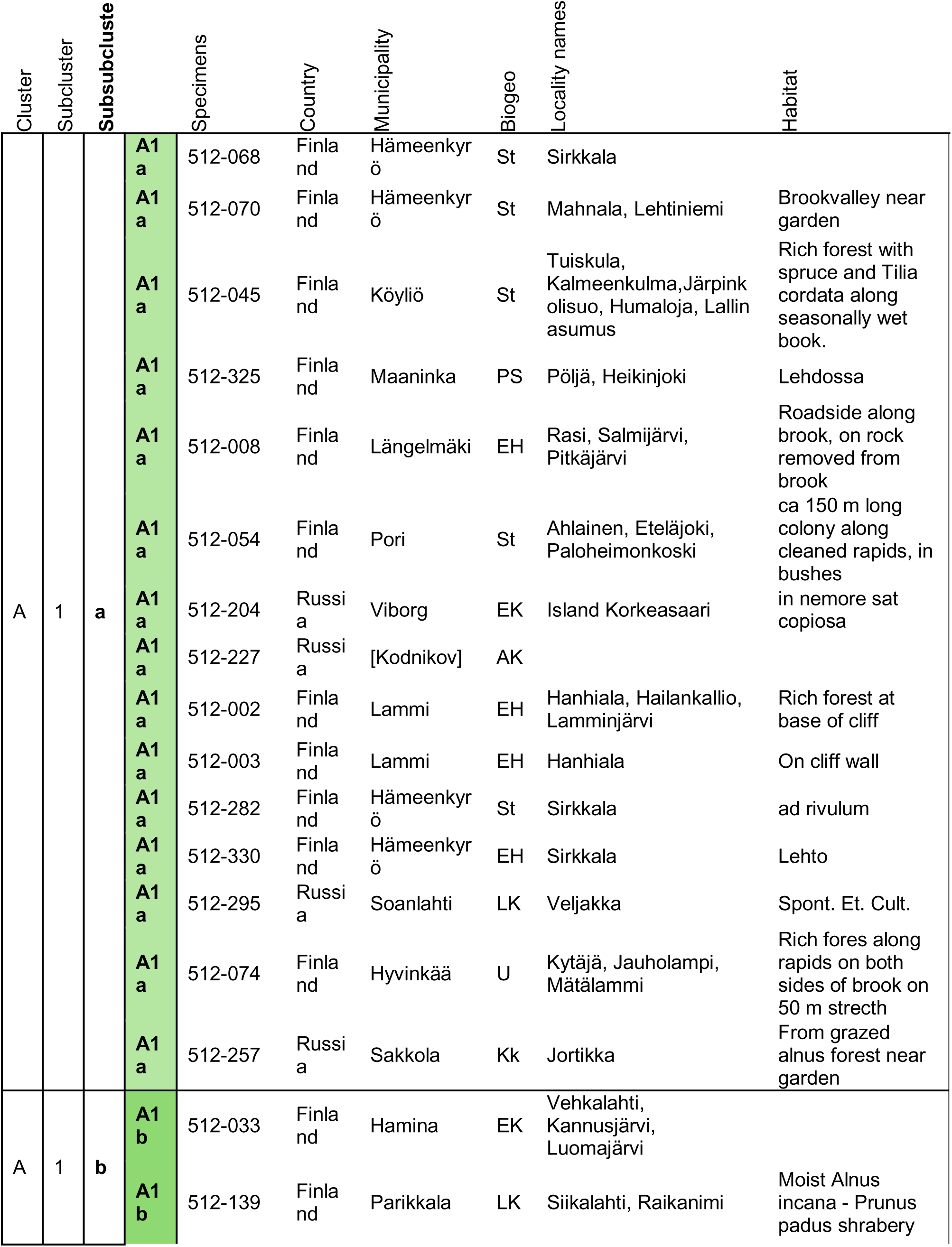
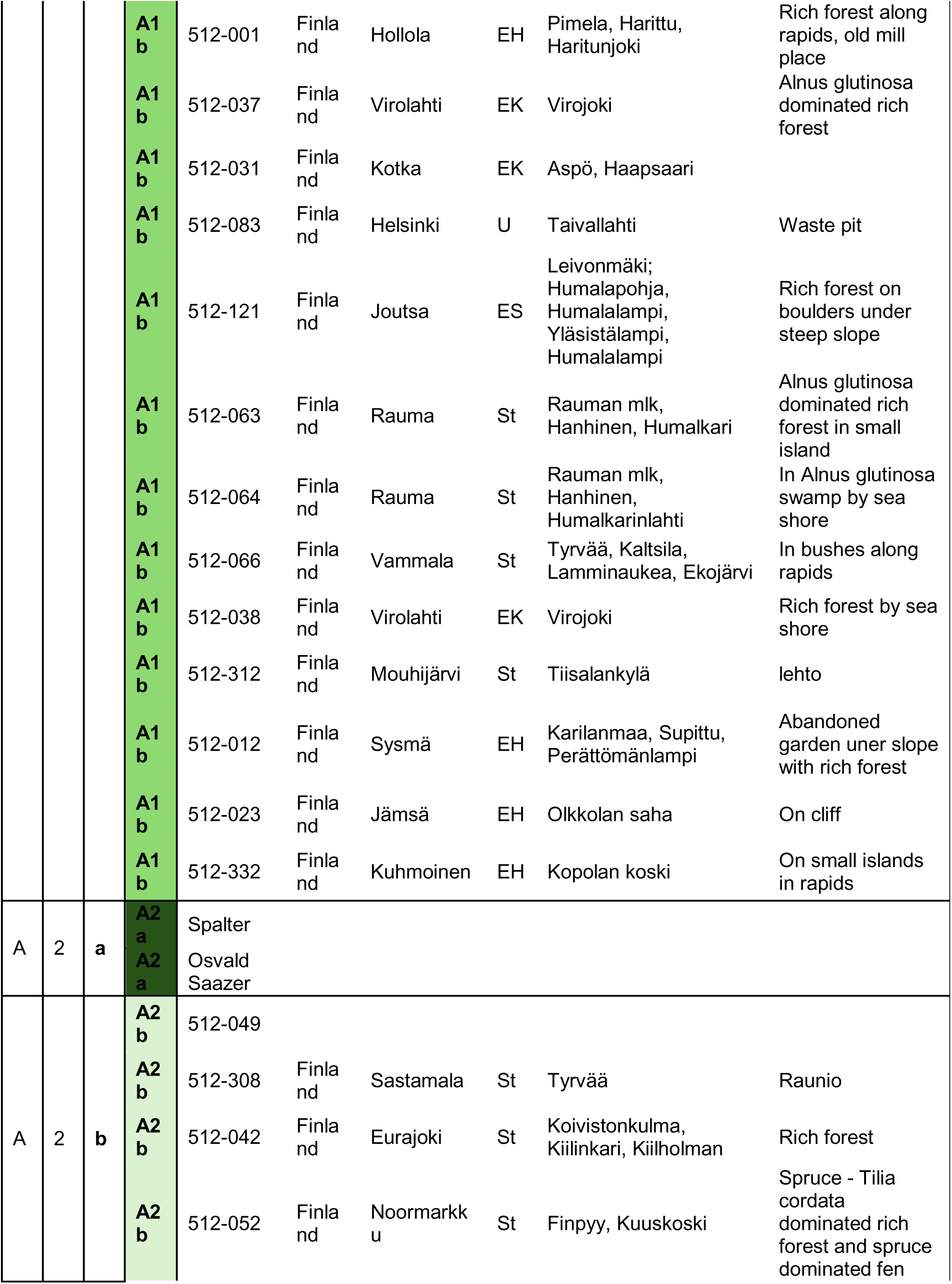
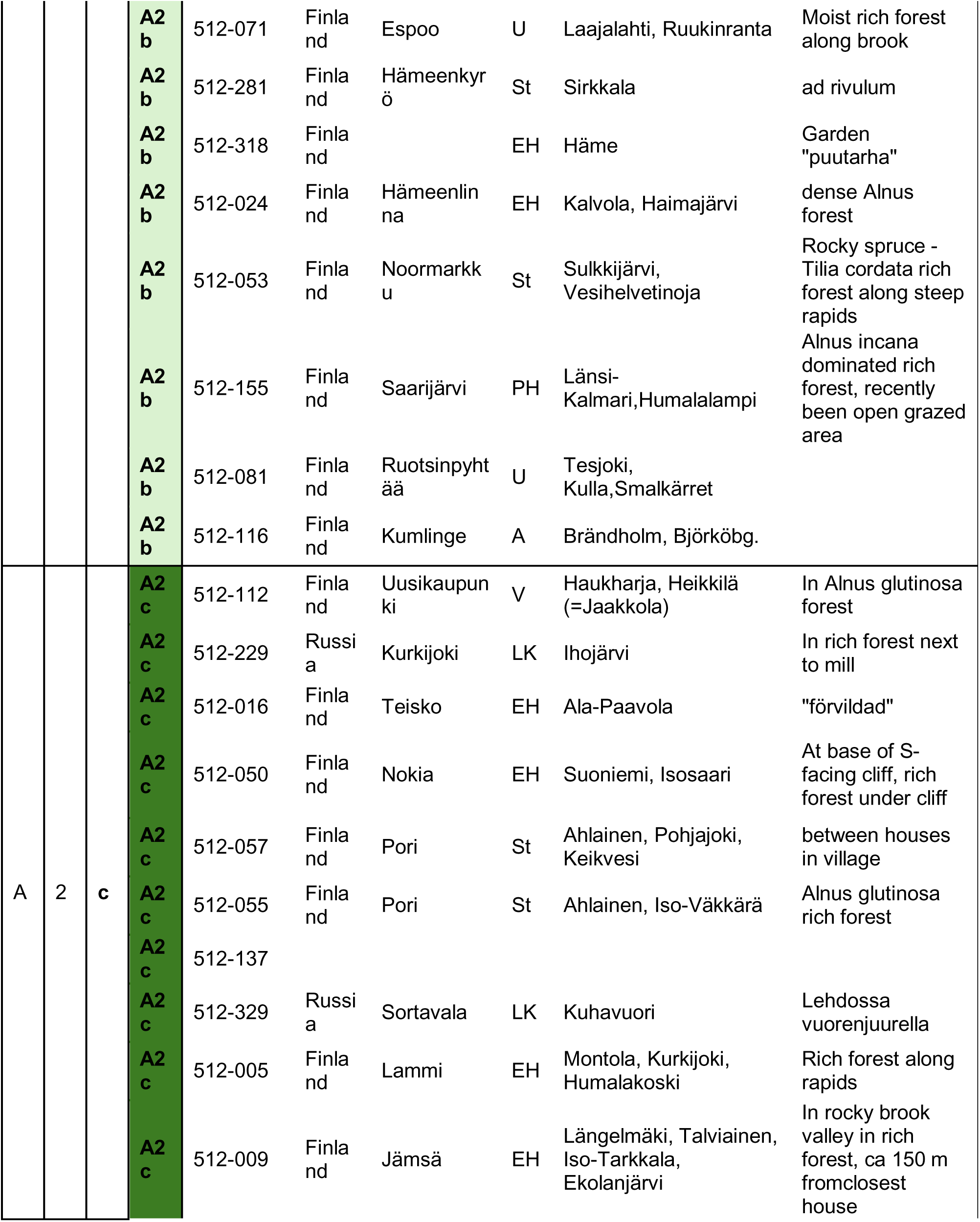
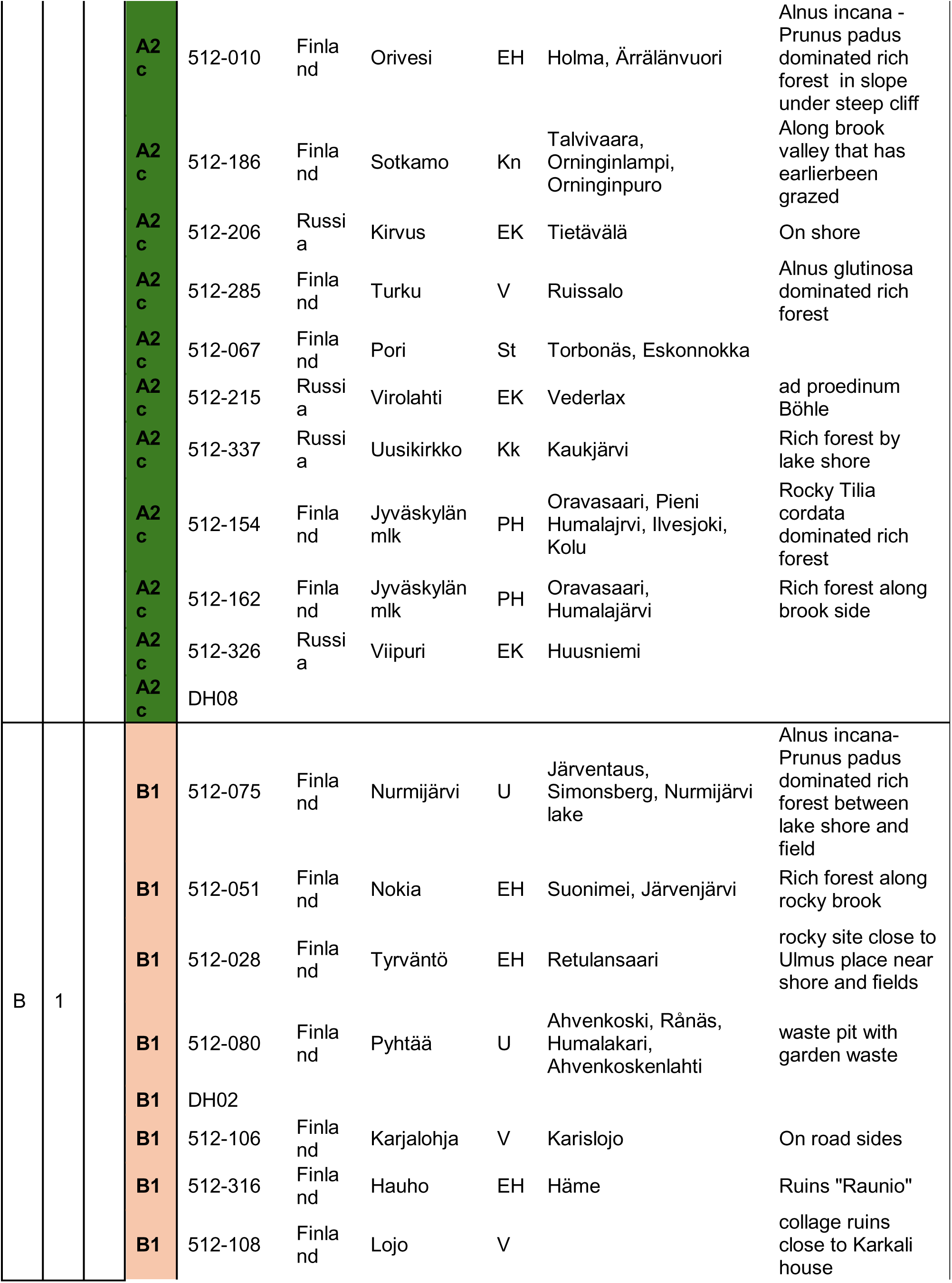
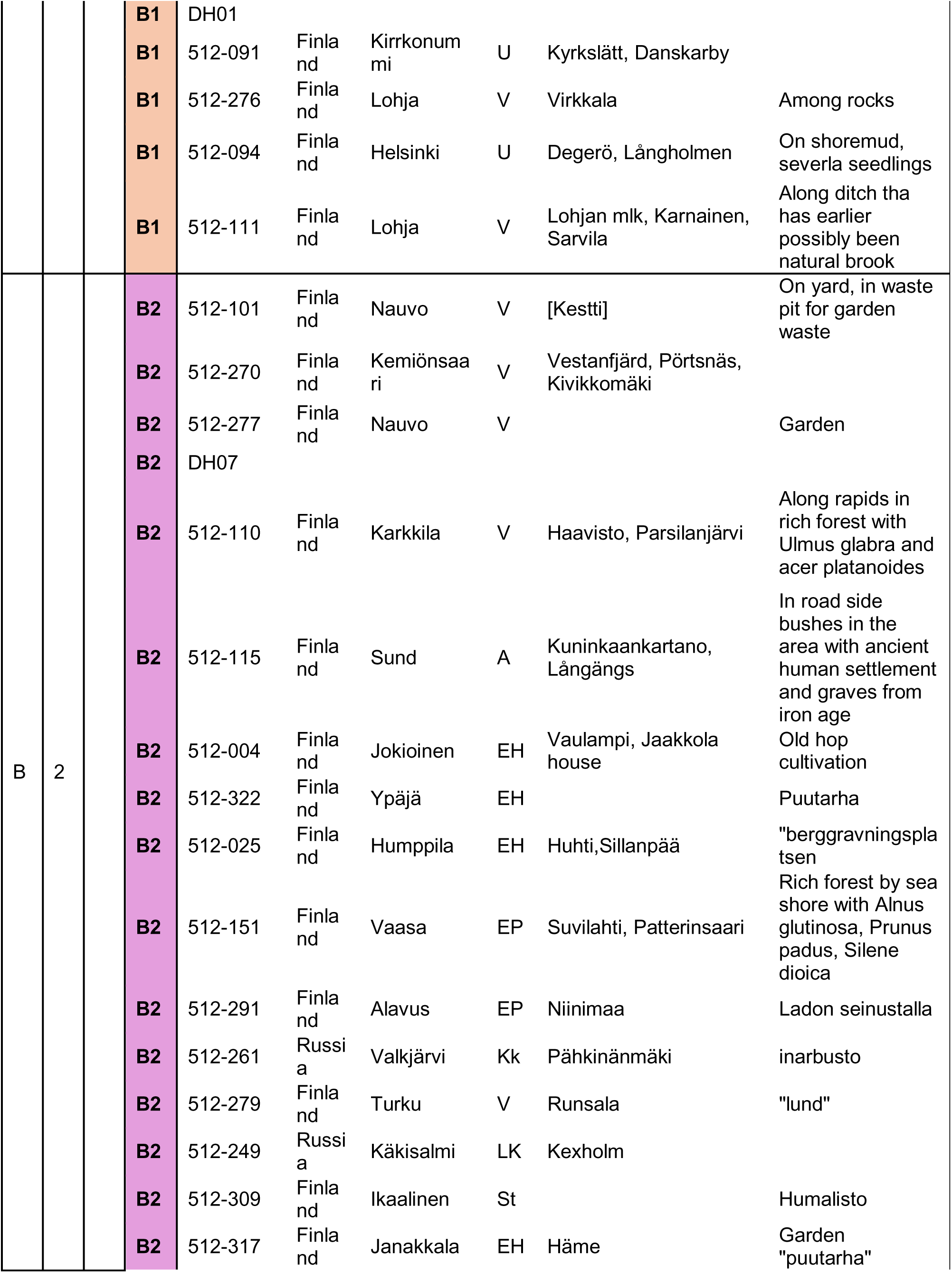
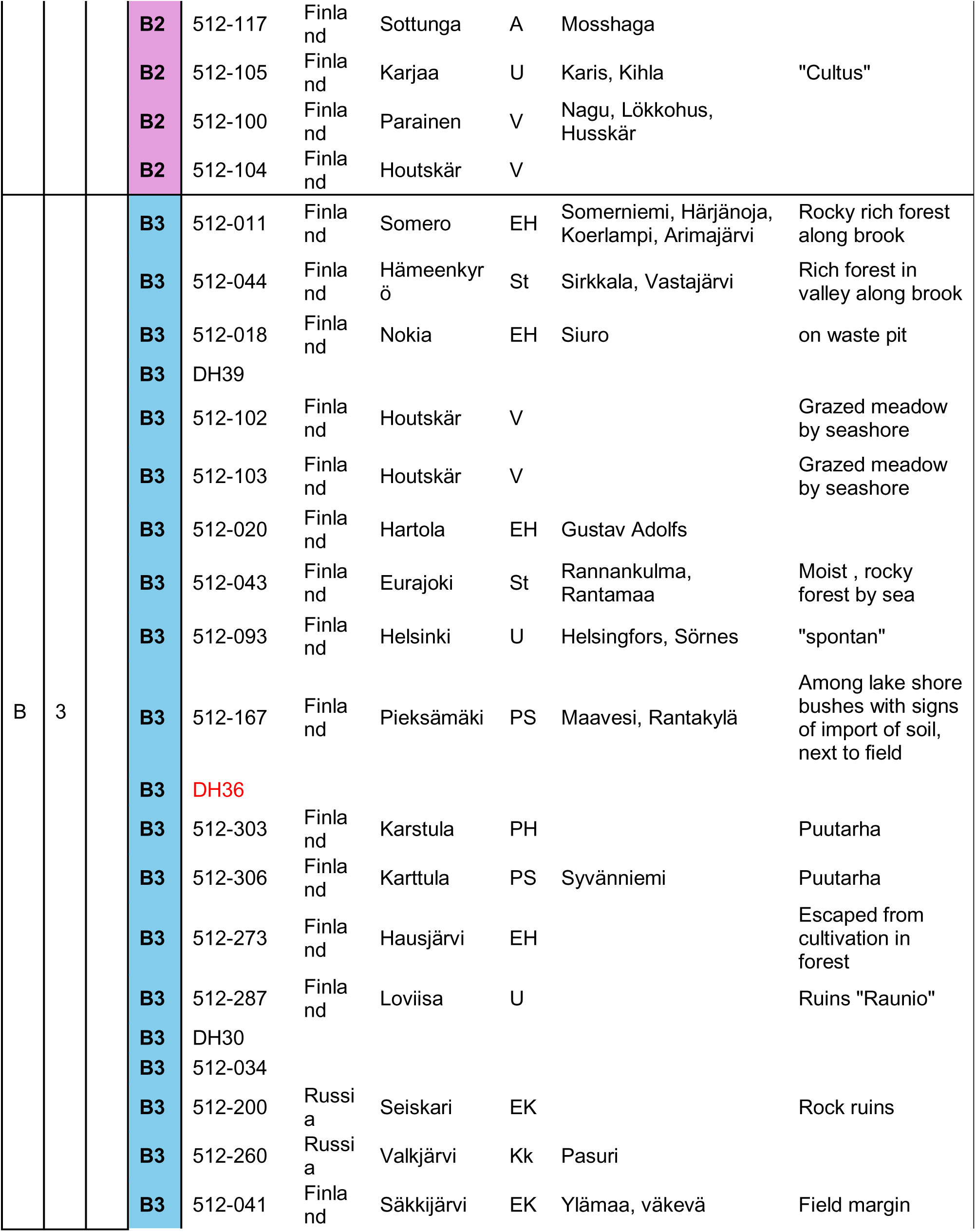
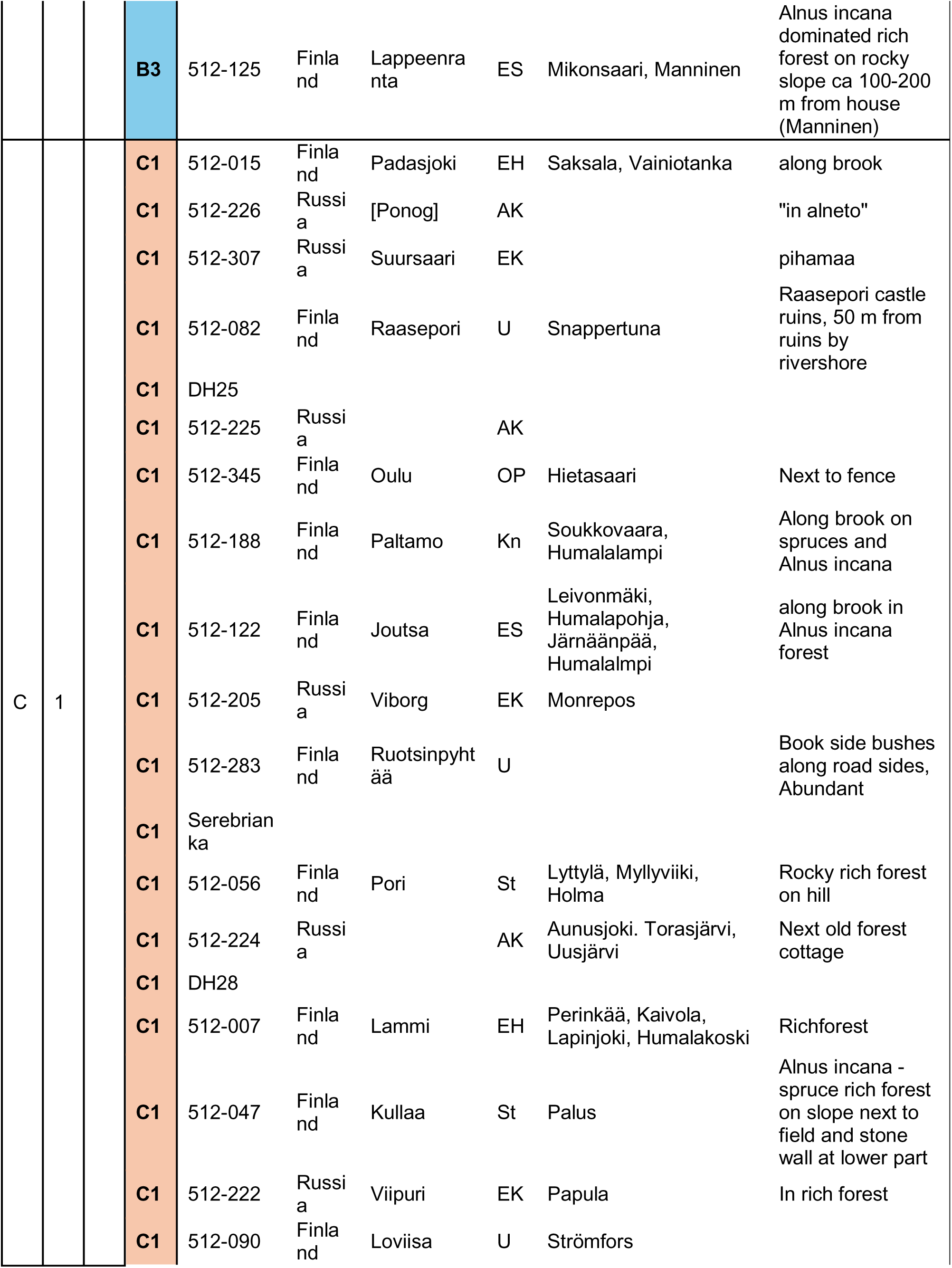
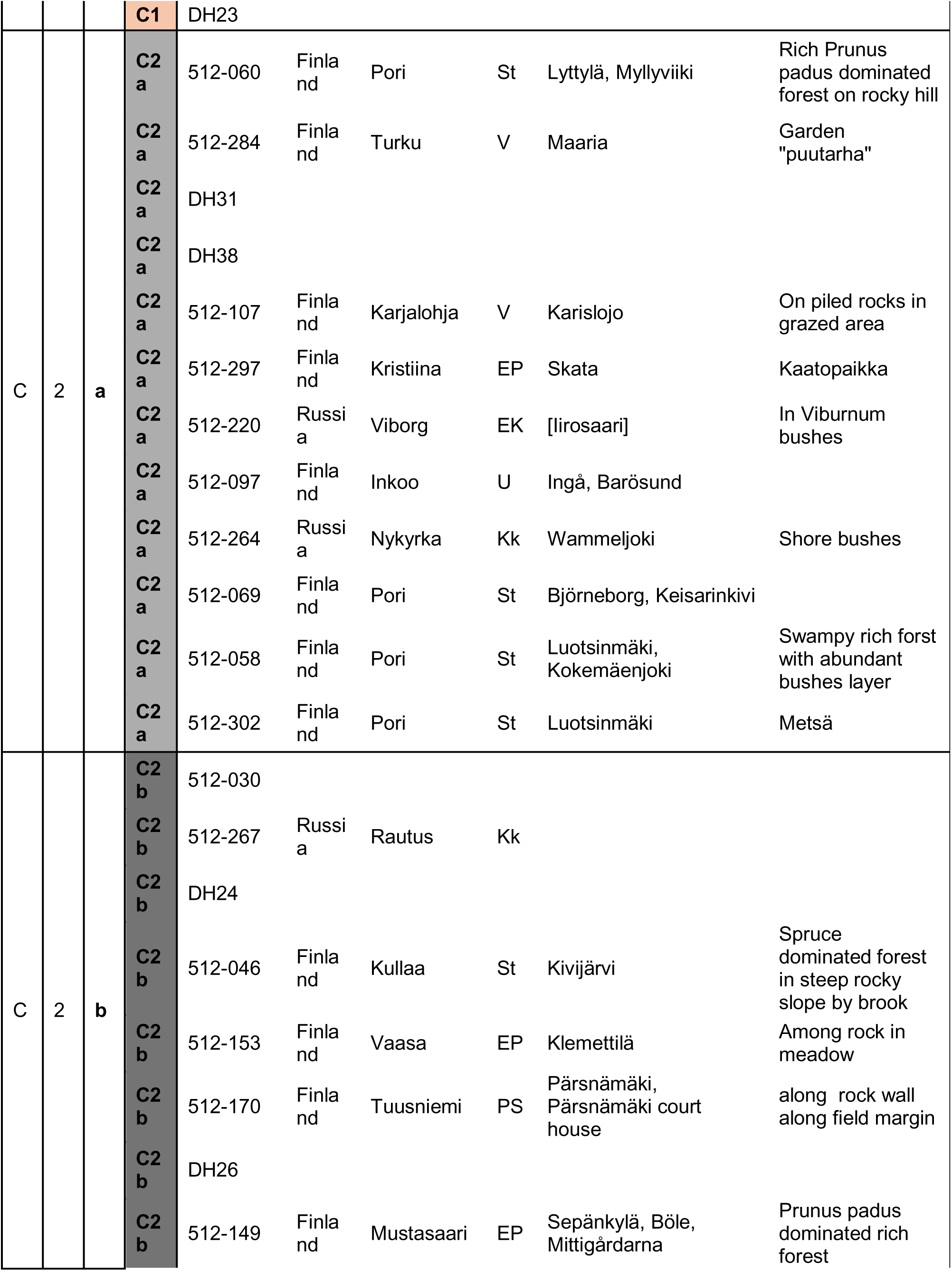
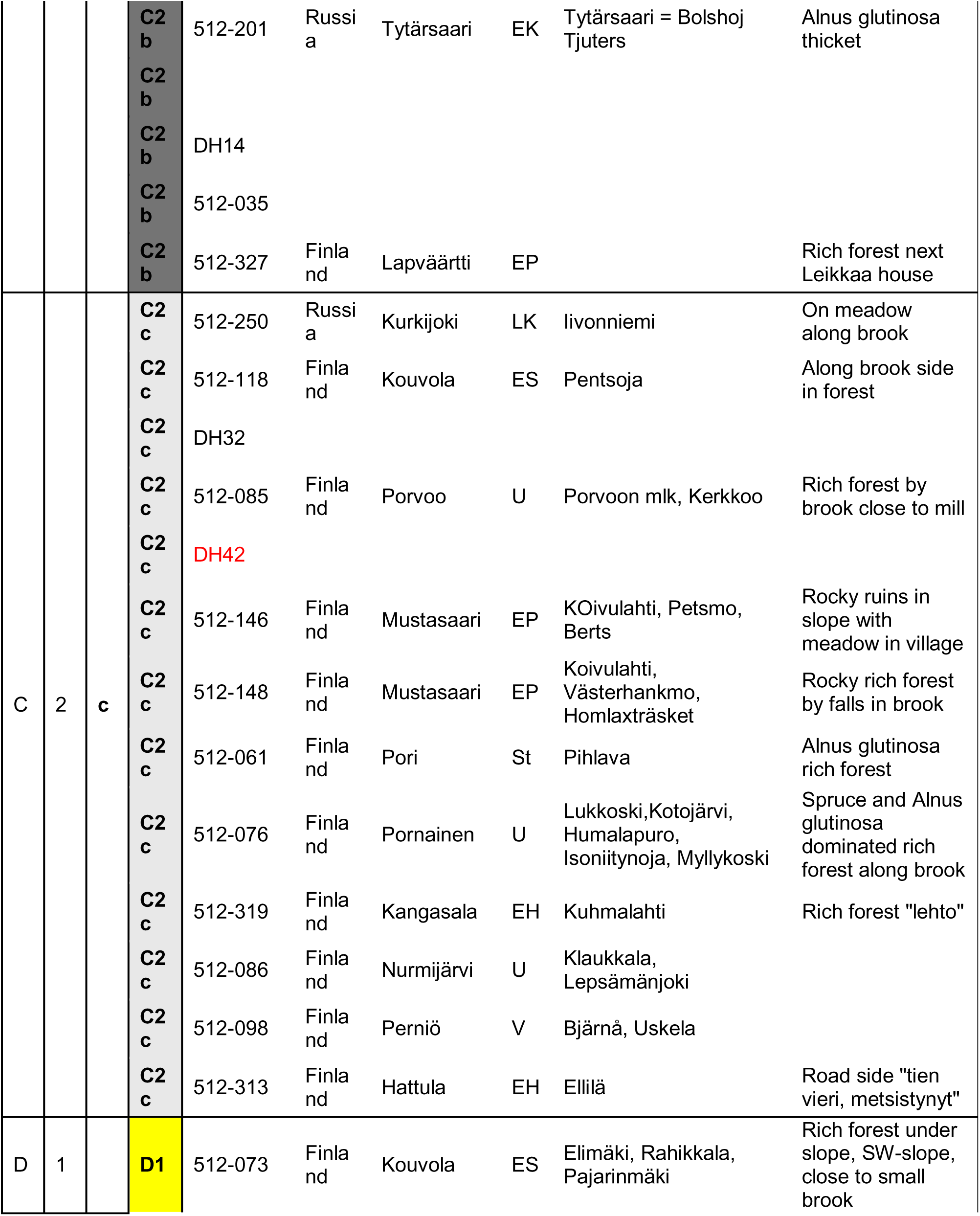
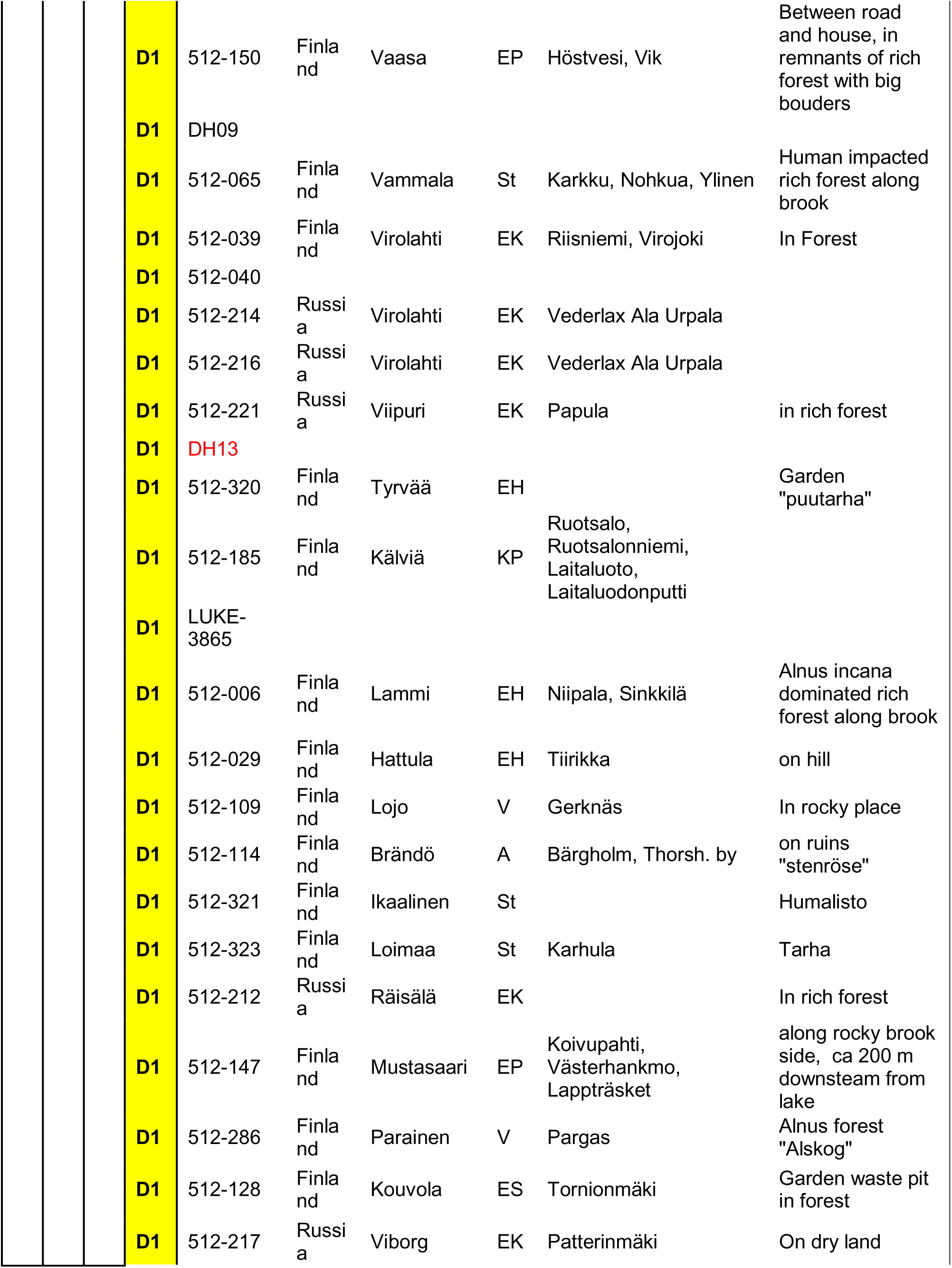
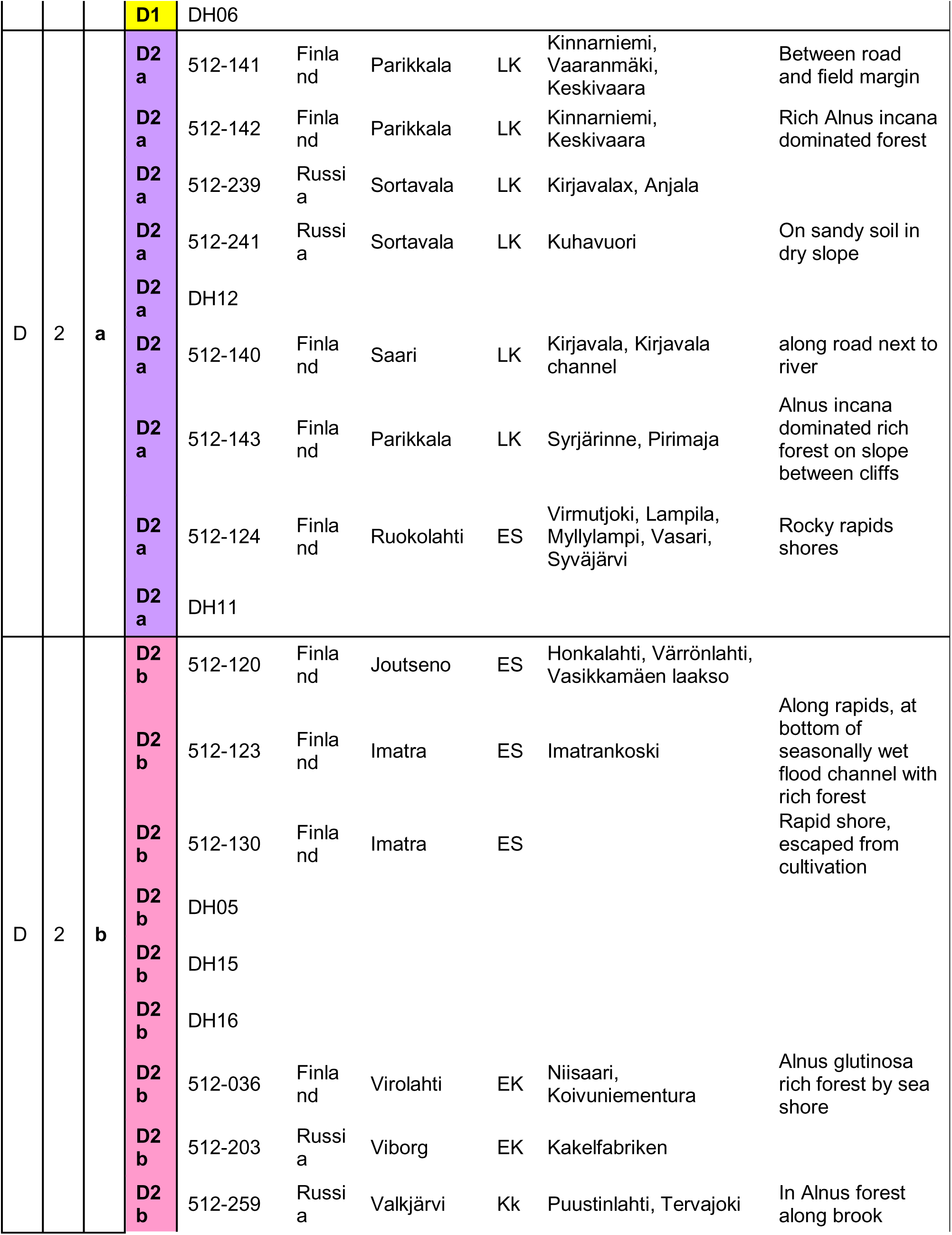
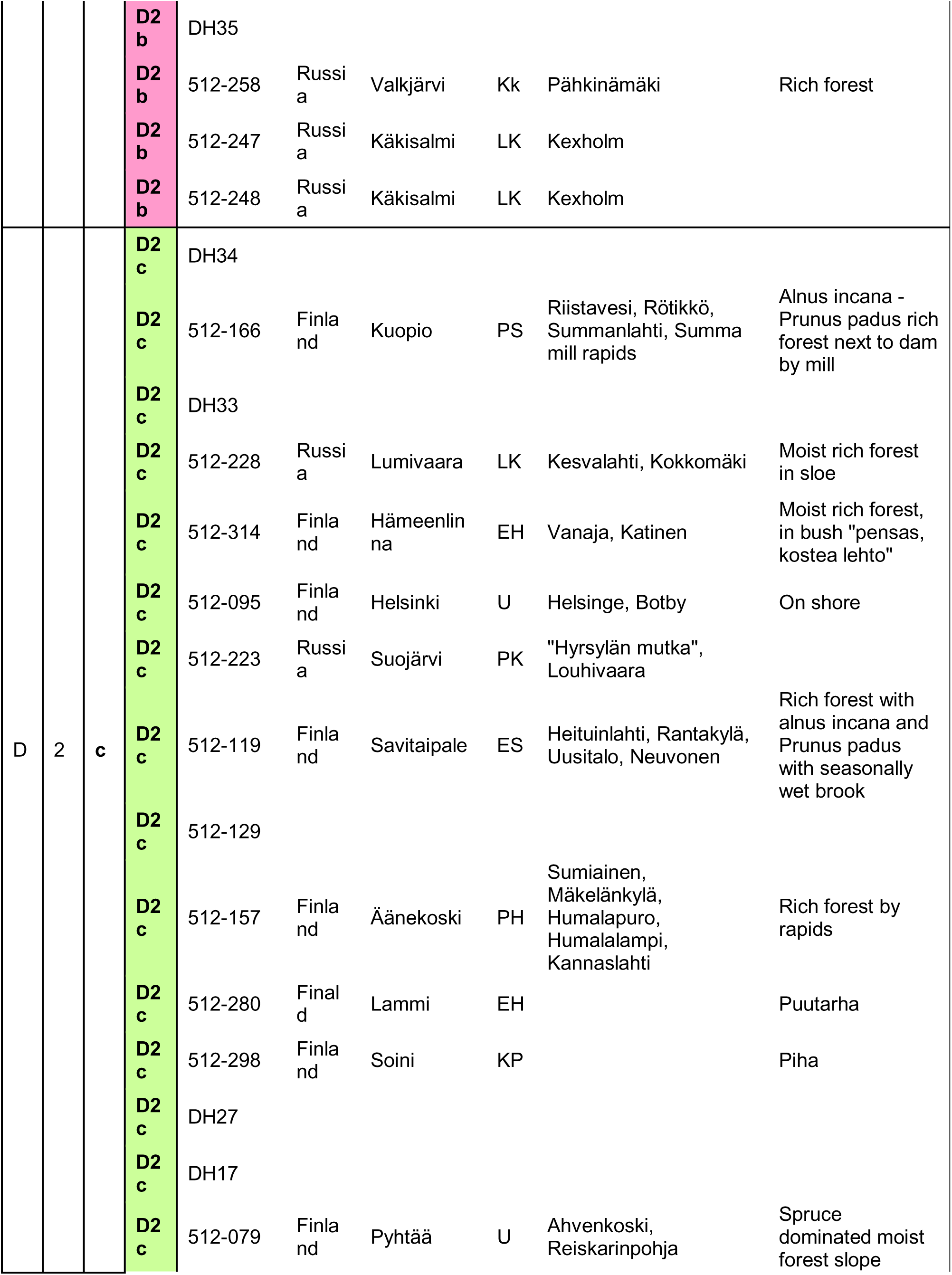
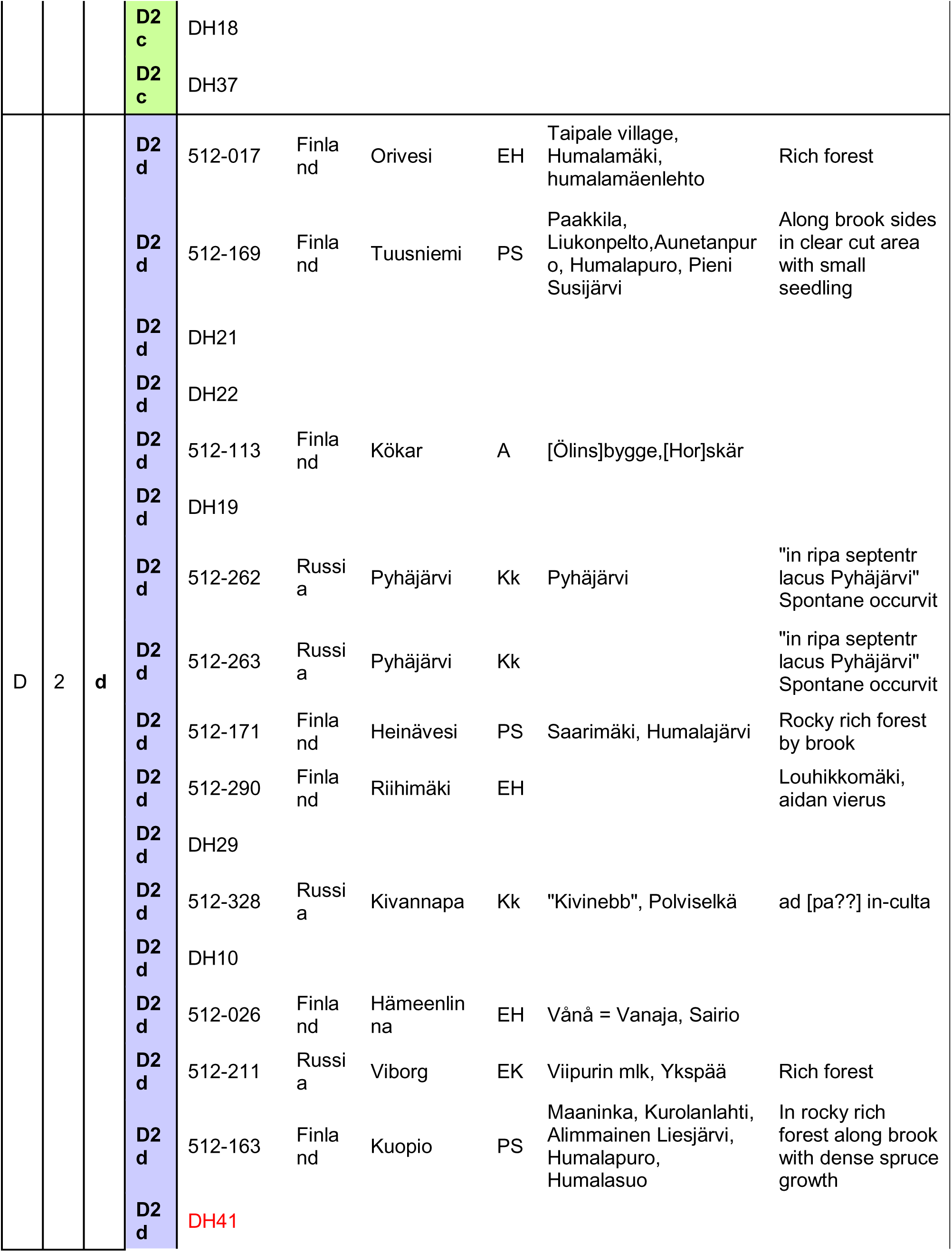
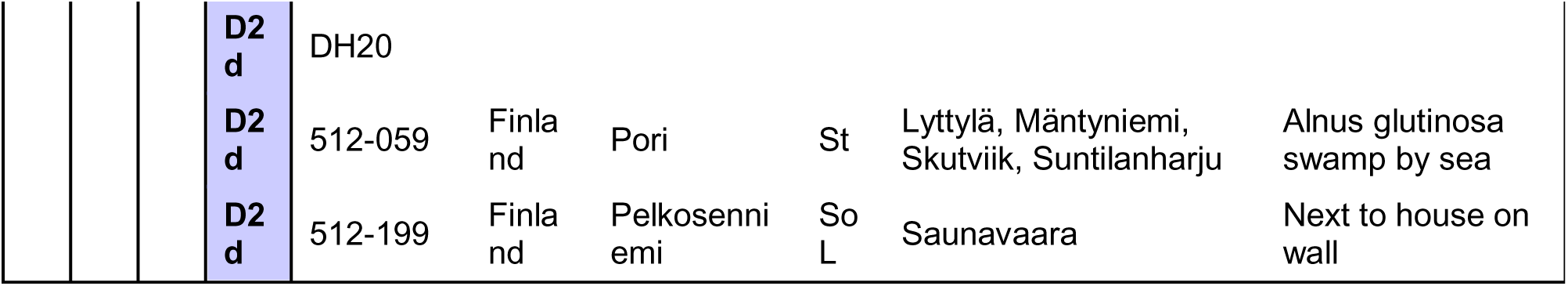

